# Kinase activity profiling identifies putative downstream targets of cGMP/PKG signaling in inherited retinal neurodegeneration

**DOI:** 10.1101/2021.09.10.459762

**Authors:** Akanksha Roy, Arianna Tolone, Riet Hilhorst, John Groten, Tushar Tomar, Francois-Paquet Durand

## Abstract

Inherited retinal diseases (IRDs) are a group of neurodegenerative disorders that lead to photoreceptor cell death and eventually blindness. IRDs are characterised by a high genetic heterogeneity, making it imperative to design mutation-independent therapies. Mutations in a number of IRD disease genes have been associated with a rise of cyclic 3’,5’-guanosine monophosphate (cGMP) levels in photoreceptors. Accordingly, the cGMP-dependent protein kinase (PKG) has emerged as a new potential target for the mutation-independent treatment of IRDs. However, the substrates of PKG and the downstream degenerative pathways triggered by its activity have yet to be determined. Here, we performed kinome activity profiling of different murine organotypic retinal explant cultures (diseased *rd1* and wild-type controls) using multiplex peptide microarrays to identify proteins whose phosphorylation was significantly altered by PKG activity. In addition, we tested the downstream effect of a known PKG inhibitor CN03 in these organotypic retina cultures. Among the PKG substrates were potassium channels belonging to the K_v_1 family (KCNA3, KCNA6), Cyclic AMP-responsive element-binding protein 1 (CREB1), DNA topoisomerase 2-α (TOP2A), 6-phosphofructo-2-kinase/fructose-2,6-biphosphatase 3 (F263), and the glutamate ionotropic receptor kainate 2 (GRIK2). The retinal expression of these PKG targets was further confirmed by immunofluorescence and could be assigned to various neuronal cell types, including photoreceptors, horizontal cells, and ganglion cells. Taken together, this study confirmed the key role of PKG in photoreceptor cell death and identified new downstream targets of cGMP/PKG signalling that will improve the understanding of the degenerative mechanisms underlying IRDs.

## Introduction

Inherited retinal degeneration (IRD) relates to a genetically highly heterogeneous group of neurodegenerative diseases causing photoreceptor cell death and eventually blindness (1,2). To this day, in almost all cases, these diseases are untreatable. Causative mutations have been identified in over 300 different disease genes (https://sph.uth.edu/retnet; information retrieved September 2021), calling for the development of mutation-independent therapies. Mutations in more than 20 IRD disease genes have been linked to increased levels of cyclic 3’,5’-guanosine monophosphate (cGMP) in photoreceptors (3) and are thought to affect at least 30% of IRD patients (4). A key effector of cGMP-signalling is cGMP-dependent protein kinase (PKG), the overactivation of which may trigger photoreceptor cell death (3,5).

Mammals possess two different genes encoding for PKG, *prkg1* and *prkg2* (6). Splicing of *prkg1* leads to two distinct isoforms – PKG1α and PKG2β – which differ in their first 80 to 100 amino acids. The *prkg2* gene gives rise to only one isoform called PKG2. PKGs exist as a homodimer and binding of cGMP to one of the four cGMP binding sites induces a conformational change which activates the kinase (6). Activated PKG phosphorylates numerous cellular proteins at serine/threonine amino acid positions, which in turn regulate numerous cellular pathways. In mammals, PKG1 regulates smooth muscle contraction (7), platelet activation and adhesion (8), cardiac function (9), feedback of the NO-signalling pathways (10), and various processes in the central nervous system, such as hippocampal and cerebellar learning (11). PKG2 is involved in translocation of CFTR channels in jejunum (12) and regulation of bone growth by activation of kinases such as MAPK3/ERK1 and MAPK1/ERK2 in mechanically stimulated osteoblasts (13).

Intriguingly, PKG also plays an important role in cell death, which has been ascertained, for instance, through studies where PKG activation inhibited tumour progression in colon cancers, breast cancers, ovarian cancers and melanoma (14,15). Furthermore, PKG seems to play a central role in photoreceptor degeneration (16–18). In the *rd1* mouse retina, a well characterised model for IRD, photoreceptor cell death is triggered by abnormally high concentrations of retinal cGMP. This event is linked to dysfunction of phosphodiesterase 6 (PDE6) – involved in the regulation of intracellular cGMP levels – caused by a nonsense mutation in the rod *Pde6b* gene (19). Increased cGMP signalling has been found in several other models for IRDs (4,16) and is likely to over-activate PKG (20). *In vitro* and *in vivo* pharmacological inhibition of PKG showed strong photoreceptor protection in *rd1* retina as well as in the retina of *rd2* and *rd10* mouse models (20,21). Together, these studies suggest a key role for PKG activity in cGMP-mediated cell death and highlight PKG as a potential common target for strategies aiming to reduce photoreceptor degeneration.

The pathways downstream of PKG in degenerating photoreceptors are nonetheless still poorly understood. Increased cGMP/PKG signalling has been associated with increased activity of poly-ADP-ribose-polymerase (PARP), histone deacetylase (HDAC), and calpain proteases, all known to be involved in photoreceptor cell death (16). However, to date there is no evidence that directly links these events to PKG. Intracellular changes of known PKG targets such as vasodilator-stimulated phosphoprotein (VASP) and cAMP response element-binding protein (CREB) have been observed in dying photoreceptors (18,22,23). While this can be a direct consequence of excessive cGMP/PKG signalling, these targets may also be phosphorylated by or have phosphorylation sites for other kinases, including cAMP-dependent protein kinase (PKA), characterised by substrate motifs similar to those of PKG (24). A better insight into the downstream effects of PKG and its phosphorylation targets is needed to understand the mechanisms of photoreceptor cell death and to, furthermore, guide the development of both new neuroprotective strategies and biomarker applications.

Using multiplex peptide microarrays, PamChips^®^, we measured PKG1- and PKG2-mediated phosphorylation of specific peptides on lysates of murine retinal explant cultures treated or not with the PKG inhibitor CN03 (21). We identified several new PKG substrates potentially connected to IRD and confirmed their retinal expression in murine tissue. This study thus provides the groundwork for future studies aimed at the elucidation of cGMP/PKG-dependent cell death pathways.

## Results

### PKG inhibition reduces photoreceptor cell death in *rd1* retinal explants

To investigate potential targets of PKG and their possible role in photoreceptor cell death, we used the PKG inhibitor CN03 on *rd1* organotypic retinal explant cultures. CN03 is a cGMP analogue, and as such is able to bind to cGMP binding sites on PKG, without inducing the conformational changes required for kinase activation (25). This culminates in reversible and competitive inhibition of PKG. In previous studies, CN03 showed marked protection of photoreceptors in retinal explants derived from the *rd1* mouse model (21,26). We therefore collected wild-type (WT) and *rd1* retinal explants treated or not with 50 μM CN03, using a treatment paradigm based on the aforementioned studies. Thus, retinas were explanted at postnatal (P) day 5 when photoreceptor degeneration had not yet started. The CN03 treatment was given at P7 and P9 and cultures were terminated at P11. The latter time-point corresponds to the beginning of *rd1* photoreceptor cell death (27), and is thus well suited to assess the protective effects of a given compound and to study events downstream of abnormal cGMP/PKG signalling. We confirmed the protective effects of CN03, by characterising the degree of cell death using the TUNEL assay (Figure 1). CN03-treated retinas showed marked photoreceptor protection as previously reported (21).

**Figure 1:**
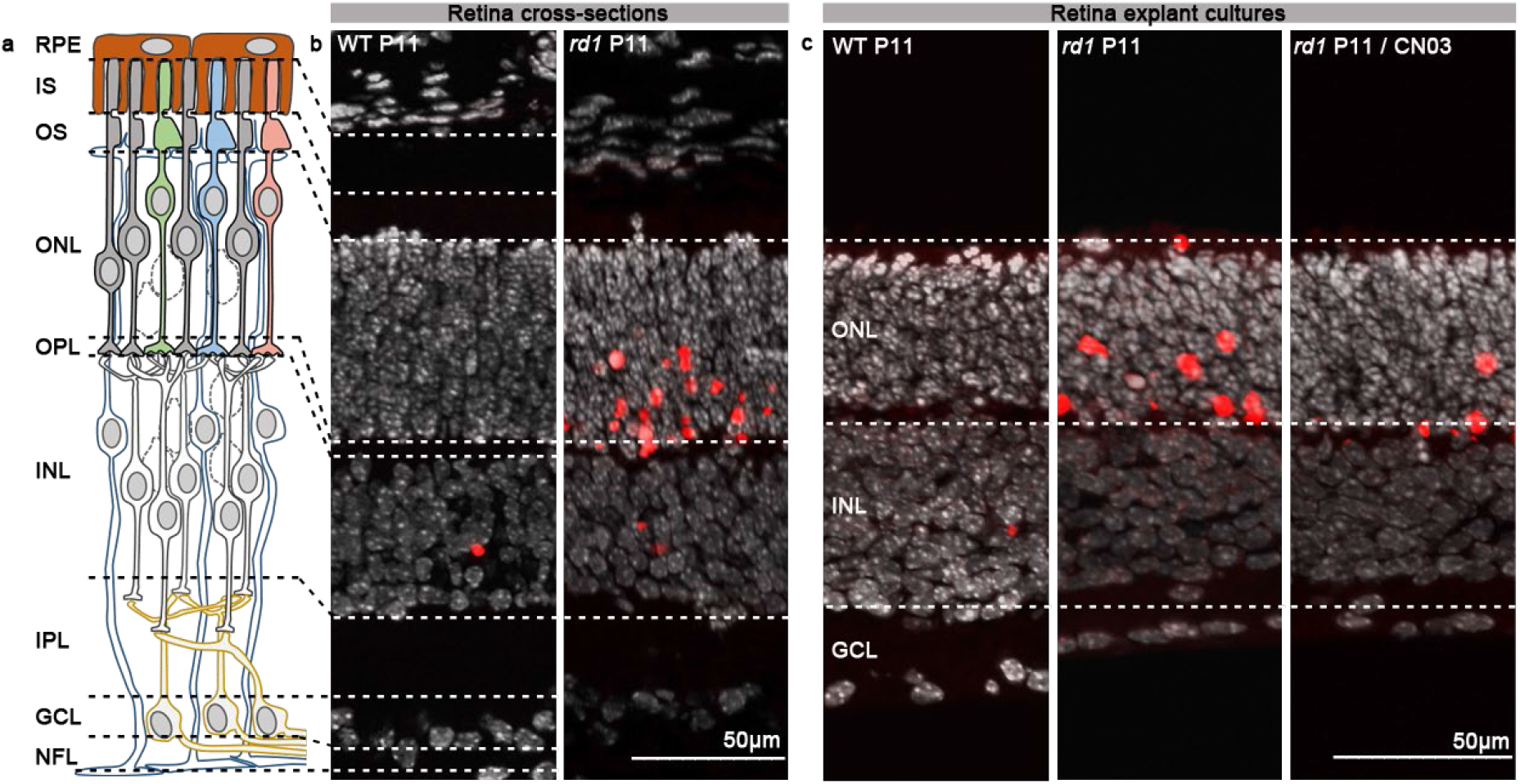
CN03-mediated photoreceptor protection in *rd1* P11 retinal explants. **a**) Diagram showing retinal layers: RPE=retina pigment epithelium, IS=inner segment, OS=outer segment, ONL=outer nuclear layer, OPL=outer plexiform layer, INL=inner nuclear layer, IPL=inner plexiform layer, GCL=ganglion cell layer, NFL=nerve fibre layer**. b**) Retina cross-sections derived from WT and *rd1* P11 mice. **c**) Sections derived from WT and *rd1* P11 retinal explant cultures untreated or treated from P7 to P11 with 50 μM CN03. In both **b** and **c**: TUNEL assay (red) indicated dying cells, DAPI (grey) was used as nuclear counterstain. P=postnatal day.

### Serine/Threonine Kinase (STK) activity in *rd1* retinal explants

To evaluate possible differences in kinase profiles of the murine retinal explant samples (*rd1*, n=8; WT, n=5), we used PamChip^®^ peptide microarray-based Serine/Threonine Kinase (STK) activity assays (24). The overall STK activity between the samples is represented as heatmap (Fig. S1 a). A violin plot was used to visualize the phosphorylation signal intensity of the peptides and its distribution within the same sample groups (Fig. 2 a). Increased phosphorylation was observed for 43% of the total 142 peptides in *rd1* retinal explants, indicating higher kinase activity in *rd1* NT when compared to WT (Fig. 2 b). Among the elevated phosphorylated peptides, SRC8_CHICK_423_435, RBL2_959_971, CDN1B_151_163 and RADI_559_569 showed significantly higher phosphorylation (*p*<0.05) in *rd1* explants than in wild type controls. Using the upstream kinase analysis tool of the BioNavigator^®^ software, the peptides with increased phosphorylation were linked to kinases that are most likely to be responsible for phosphorylation of these peptides (See Data Analysis, Materials and Methods Section). The kinase statistics and kinase score were calculated as a metrics for identifying highly active kinases. The kinases that were predicted to be highly active in *rd1* explants include CaMK4, PKG1, PKG2, PKAα, and Pim1 (Table 1). In order to present relative kinome activity profiles of retinal explants (*rd1* vs WT), kinase score and kinase statistics were used to visualize branches and nodes on the phylogenetic tree of protein kinase family (Fig. 2 c).

**Table 1.**
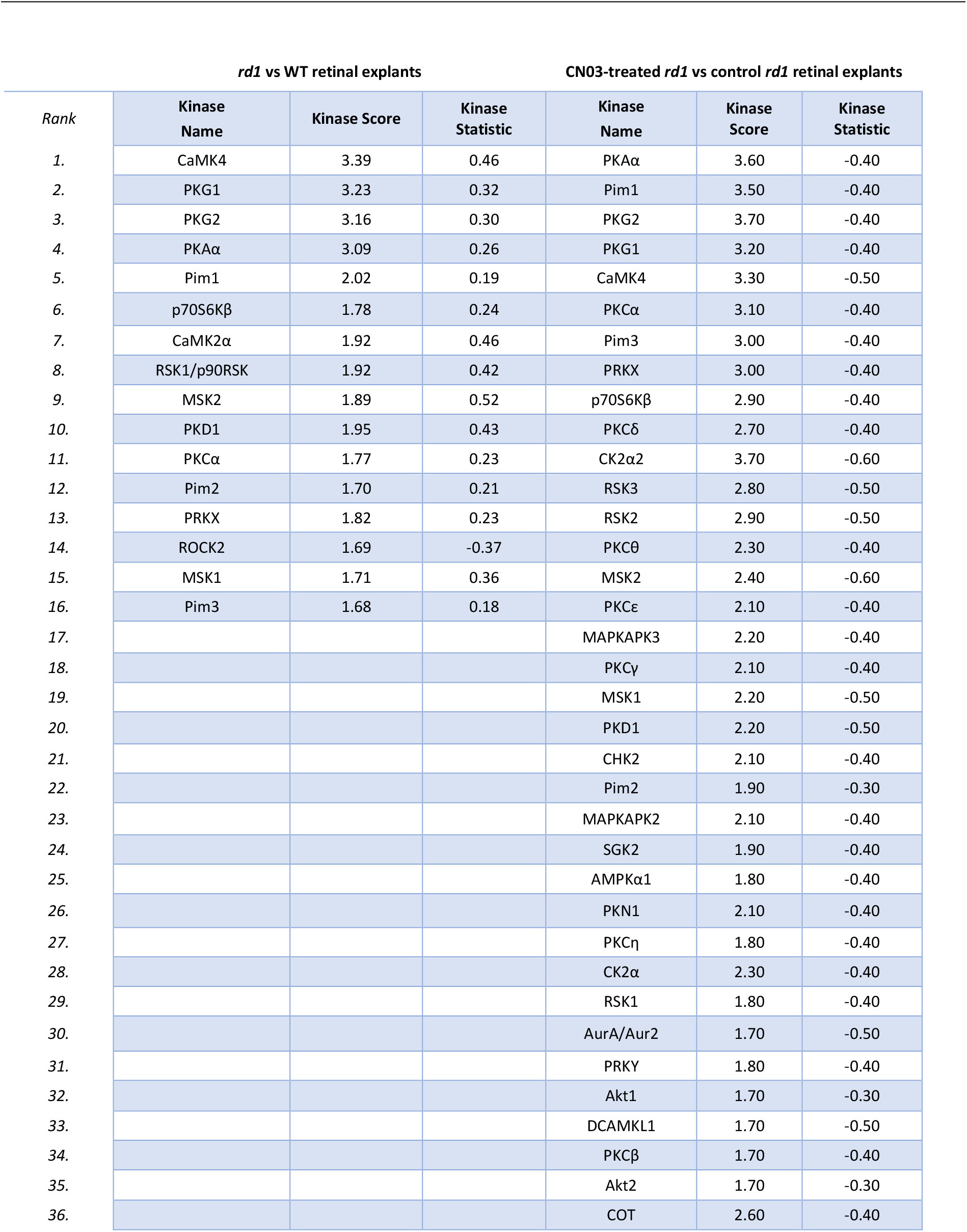
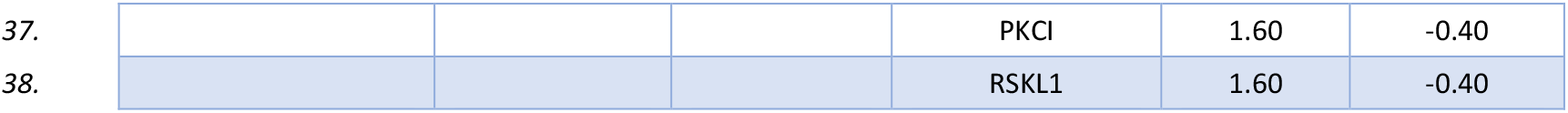
Upstream kinase analysis results for *rd1 vs*. WT. **(a)** and *rd1* CN03 *vs*. *rd1* **(b)**. The kinase statistic shows the overall change of the peptide set that represents a given kinase. Positive values indicate higher kinase activity in *rd1*, while negative value indicate lower activity in *rd1* CN03. The kinase score includes the sum of significance and specificity score (Score greater than 1.5 shown in the table). This score indicates the significance of a change represented by the kinase statistic. The higher the score, the higher the significance of the change. The specificity score indicates the specificity of the kinase statistic in terms of the set of peptides used for the corresponding kinase. The higher the score, the less likely it is that observed kinase statistic could have been obtained using a random set of peptides from the data set.

**Figure 2.**
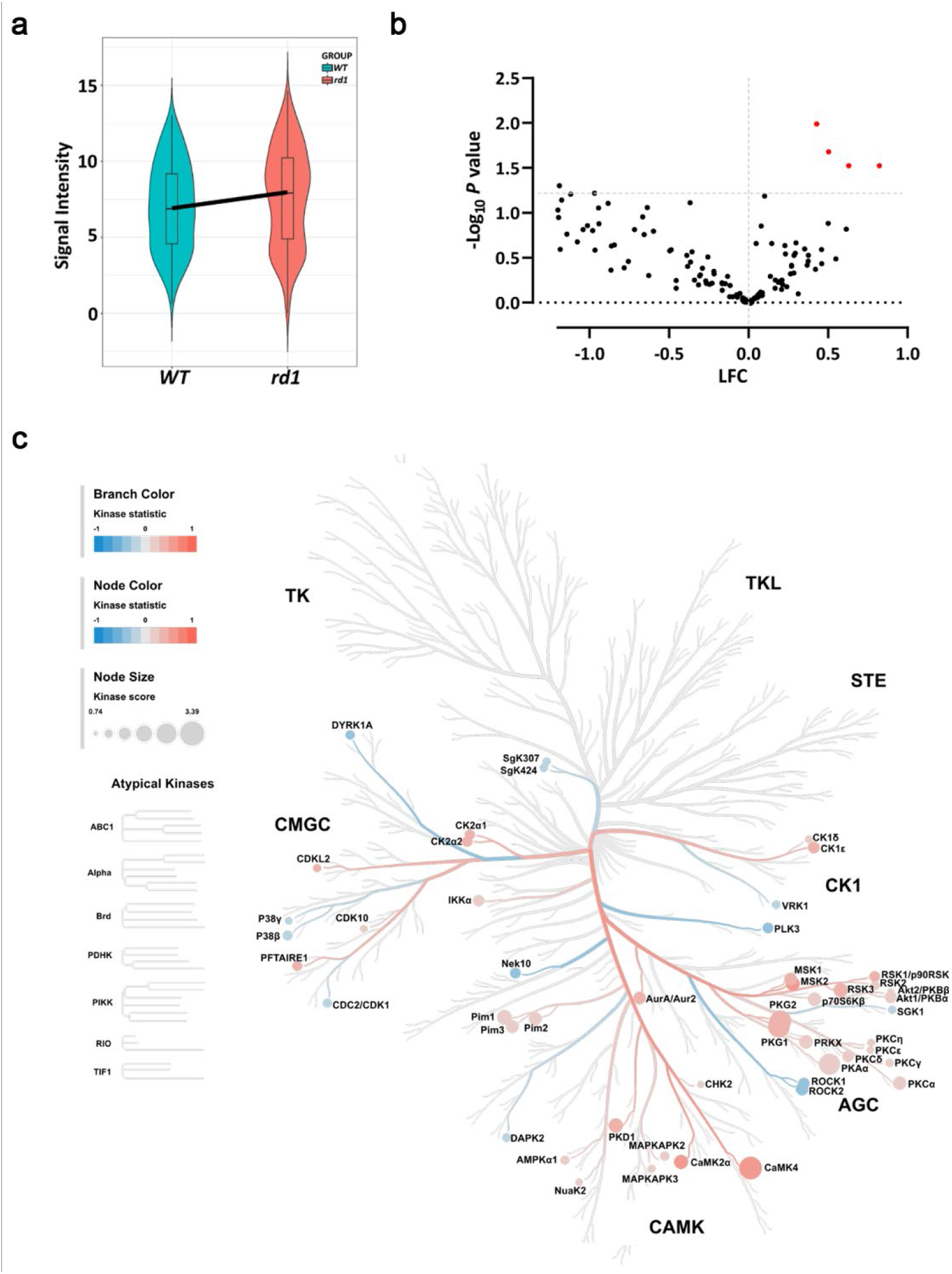
Serine/Threonine Kinase (STK) activity in untreated retinal explants. Organotypic retinal explants derived from wild-type (WT) and *rd1* mice (WT, n=5; *rd1*, n=8) were treated with culture medium for four days. The kinase activity of their lysates was measured on PamChip^®^ Serine/Threonine kinase (STK) arrays. **a**) Violin plot showing the global phosphorylation of the peptides on PamChip^®^ STK array as Log_2_ signal intensity and their intensity value distribution, when comparing WT to *rd1* explants. The thick line connects the average phosphorylation values of each group. **b**) Volcano Plot representing Log Fold Change (LFC) and -Log_10_ *p-*value for peptide phosphorylation. Red dots indicate significantly changed phosphopeptides (*p-*value < 0.05), black dots represent peptides with no significant alteration in phosphorylation. **c**) The high-ranking kinases were visualized in a kinome phylogenetic tree, where branch and node color are encoded according to the kinase statistic value with value > 0 (in red) representing higher kinase activity in *rd1* retinal explants. The kinase score is encoded in node size where kinases were ranked based on their significance and specificity in terms of set of peptides used for the corresponding kinase.

### STK activity in *rd1* retinal explants treated with PKG Inhibitor CN03

As we found a higher phosphorylation of peptides in *rd1* explants, when compared to WT explants, and with a clear role for PKGs observed, we sought to investigate the effect of the PKG inhibitor CN03 on the STK activity. The PKG inhibitor CN03 significantly decreased photoreceptor cell death (*cf*. Fig. 1). The overall STK activity profiles for treated and untreated *rd1* retinal explants are shown in a heatmap (Fig. S1 b). The distribution of phosphorylated peptides for both the samples is represented by a Violin plot (Fig. 3 a). Phosphorylation decreased for approximately 80% of the 142 peptides present on the STK PamChip^®^. Fourteen peptides were identified whose phosphorylation decreased significantly (*p*<0.05) in *rd1* CN03 as compared to untreated *rd1* (Fig. 3 b). Table 2 shows peptides that displayed lower phosphorylation (22 peptides, *p*<0.1) in CN03 treated *rd1* retina than untreated controls with their UniProt IDs, names of the proteins they are derived from, *p*-value, substrate preference of PKG1 and PKG2, and localization within the retina.

**Table 2.** List of potential PKG targets. Peptide substrates for PKG1and PKG2 that were differentially phosphorylated in *rd1* untreated *vs*. *rd1* CN03 treated retinal explant cultures. The table shows the peptide name, UniProt ID, Protein name, *p* value, PKG1 and PKG2 substrate specificity score, and localization in the retina with reference. The scores for PKGI, PKGII, and PKA range from 1 to 10. For substrates previously detected in the mouse retina, the reference is added. The expression of substrates printed in bold was further studied using immunodetection (*cf*. Fig. 4). Colour scheme according to the PKG specificity score: 10–8: Good, 7–4: Intermediate and 3–0: Poor substrate (24). For substrates known to be present in the mouse retina, the reference is added. The expression of substrates printed in bold was further studied using immunodetection (*cf*. Fig. 4).

| S. No. | ID | UniProt ID | Protein name | p value | PKG1 score | PKG2 score | Localisation | Ref. |
| --- | --- | --- | --- | --- | --- | --- | --- | --- |
| 1. | IKBA_26_38 | P25963 | NF-kappa-B inhibitor alpha | 0.016 | 0 | 0 | ONL, INL, GCL | (28) |
| 2. | GBRB2_427_439 | P47870 | Gamma-aminobutyric acid receptor subunit beta-2 precursor | 0.026 | 10 | 6 |  |  |
| 3. | PTN12_32_44 | Q05209 | Tyrosine-protein phosphatase non-receptor type 12 | 0.031 | 10 | 5 |  |  |
| 4. | KCNA6_504_516 | **P17658** | **Potassium voltage-gated channel subfamily A member 6** | **0.032** | 9 | 3 | GCL | (29) |
| 5. | NCF1_296_308 | P14598 | Neutrophil cytosol factor 1 | 0.032 | 9 | 7 | GCL | (30) |
| 6. | CREB1_126_138 | **P16220** | **cAMP response element-binding protein** | **0.034** | 10 | 6 | ONL, INL, GCL |  |
| 7. | GRIK2_708_720 | **Q13002** | **Glutamate receptor, ionotropic kainate 2 precursor** | **0.035** | 10 | 4 | INL, IPL | (31) |
| 8. | NCF1_321_333 | P14598 | Neutrophil cytosol factor 1 | 0.036 | 0 | 0 |  |  |
| 9. | EPB42_241_253 | P16452 | Erythrocyte membrane protein band 4.2 | 0.037 | 9 | 7 |  |  |
| 10. | VTNC_390_402 | P04004 | Vitronectin precursor | 0.037 | 10 | 4 | RPE, ONL, GCL | (32) |
| 11. | ADRB2_338_350 | P07550 | Beta-2 adrenergic receptor | 0.039 | 8 | 4 | Müller cells | (33) |
| 12. | BRCA1_1451_1463 | P38398 | Breast cancer type 1 susceptibility protein | 0.039 | 0 | 0 |  | (34) |
| 13. | TOP2A_1463_1475 | **P11388** | **DNA topoisomerase 2-alpha** | **0.039** | 6 | 5 | GCL |  |
| 14. | MPIP3_208_220 | P30307 | M-phase inducer phosphatase 3 | 0.041 | 0 | 0 |  |  |
| 15. | CDC2_154_169 | P06493 | Cell division control protein 2 homolog | 0.046 | 0 | 0 |  |  |
| 16. | KAP3_107_119 | P31323 | cAMP-dependent protein kinase type II-beta regulatory subunit | 0.046 | 8 | 4 |  |  |
| 17. | MYPC3_268_280 | Q14896 | Myosin-binding protein C, cardiac-type | 0.049 | 10 | 4 |  |  |
| 18. | RYR1_4317_4329 | P21817 | Ryanodine receptor 1 | 0.053 | 9 | 5 | ONL, INL, GCL | (35) |
| 19. | TY3H_65_77 | P07101 | Tyrosine 3-monooxygenase | 0.054 | 9 | 3 |  |  |
| 20. | KCNA3_461_473 | **P22001** | **Potassium voltage-gated channel subfamily A member 3** | **0.055** | 8 | 7 | ONL, IPL | (29) |
| 21. | F263_454_466 | **Q16875** | **6-phosphofructo-2-kinase/fructose-2,6-biphosphatase 3** | **0.059** | 10 | 3 | INL, GCL |  |
| 22. | VASP_150_162 | P50552 | Vasodilator-stimulated phosphoprotein | 0.081 | 8 | 4 | ONL | (23) |

**Figure 3.**
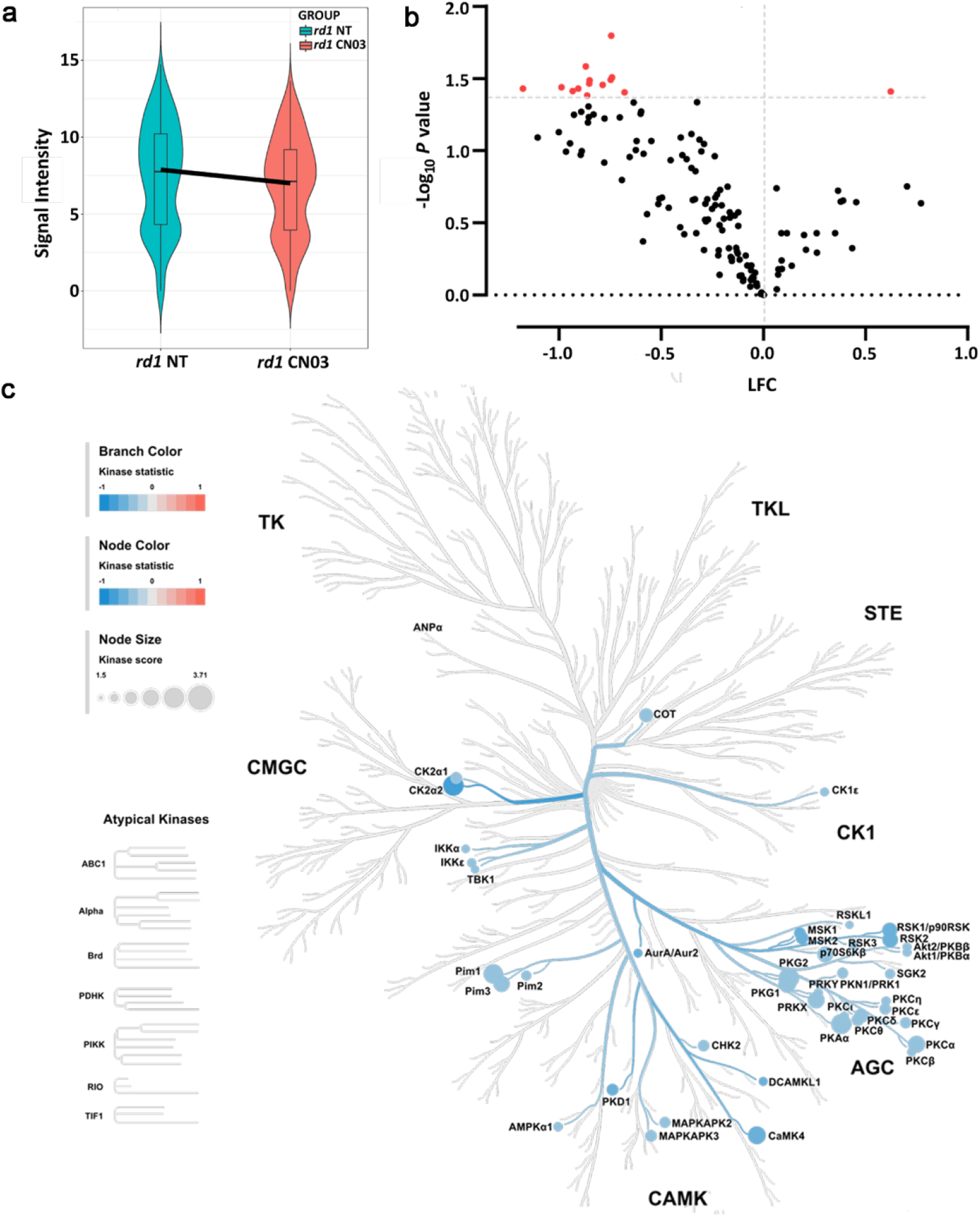
Serine/Threonine Kinase (STK) activity in response to PKG inhibition in retinal explants. Retinal explant cultures were either non-treated (NT) or treated with 50 μM CN03 for 4 days (*rd1* NT, n=8; *rd1* CN003, n=10). The kinase activity of retinal explant lysates was measured on PamChip^®^ Serine/Threonine kinase (STK) arrays. **a**) Violin plot showing the global phosphorylation of peptides on the PamChip^®^ STK array as Log_2_ signal intensity and their intensity value distribution, when comparing *rd1* NT to *rd1* treated CN03 explants. The thick line is connecting the average phosphorylation values of each group. **b**) Volcano Plot representing Log Fold Change (LFC) and -Log_10_ *p* value) for peptide phosphorylation. Red dots indicate significantly changed phosphopeptides with *p* value < 0.05 and black dots represent phosphopeptides with no significant alteration in phosphorylation. **c**) The high-ranking kinases are visualized in a kinome phylogenetic Tree, where branch and node color are encoded according to the kinase statistic value with value < 0 (in blue) representing lower kinase activity in *rd1* retinal explants treated with CN03. The kinase score is encoded in node size where kinases were ranked based on their significance and specificity in terms of set of peptides used for the corresponding kinase.

**Figure 4.**
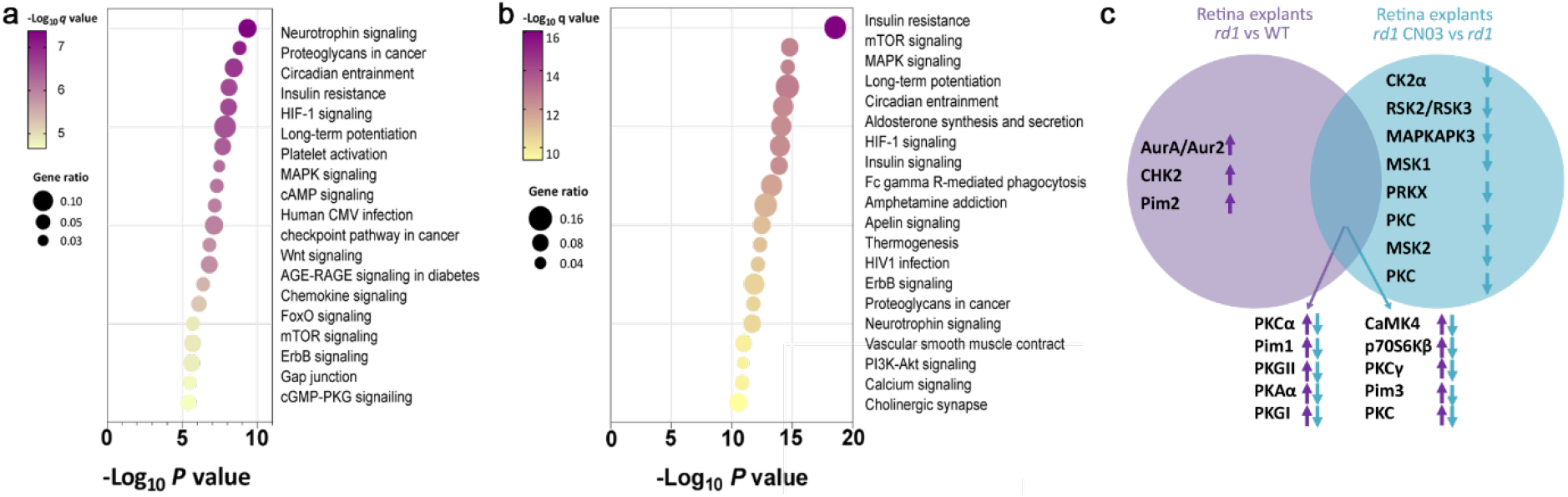
Biological pathways involved in retinal degeneration. Key biological pathways potentially higher activity in *rd1* **(a)** and lower activity in *rd1* treated with CN03 **(b)**. Pathways are ranked according to their *p-* values and colored by their *q*-values. A *q* value is the *p*-value adjusted for multiple testing using the Benjamini-Hochberg procedure (36,37). The node size indicates the gene ratio, *i.e*., the percentage of total genes or proteins in the given KEGG pathways (only input genes or proteins with at least one KEGG pathway were included in the calculation). Node color represents the false discovery rate. **c)** The overlap between kinases changed in *rd1* NT vs WT and vs *rd1* treated with CN03 respectively, is visualized in a Venn Diagram. Here, increased or reduced activity of a kinase is indicated by arrows pointing up- or down-wards, respectively.

The designation of the peptides as PKG1- or PKG2-specific substrates is based on our recent studies where we ranked the peptides on the STK PamChip^®^ according to their preference for PKG1 and or PKG2 (24). This preference was based on the substrate phosphorylation in response to PKG activity modulators (ATP, cGMP, cAMP, PKG activator, PKG inhibitors), in the presence of recombinant PKG1 or PKGII. In the present study, 11 of the peptides in Table 2, are substrates for both PKG1 and PKG2 and one was for PKG1 only. Among those peptides, KCNA6_504_516, NCF1_296_308, GRIK2_708_720, VTNC_390_402, ADRB2_338_350, BRCA1_1451_1463, RYR1_4317_4329, KCNA3_461_473 and VASP_150_162 have been verified to be present in the retina by literature search and/or ex vivo analysis.

We then linked the phosphorylated peptides to the putative upstream kinases and found that kinase activity of particularly PKAα, Pim1, PKG1, PKG2, CaMK4 was suggested to be reduced by CN03 treatment (Table 1, Fig. 3 c,). Notably, these were the same kinases that were predicted to be more active in the diseased *rd1* explants in comparison to WT explants, indicating specific targeting by CN03 (Fig. 2 c).

### Putative biological pathways involved in retinal degeneration

To identify possible associations of the kinases with biological pathways that are activated in retinal degeneration as represented by *rd1* explants, we performed an analysis of relevant biochemical pathways using the Kyoto Encyclopedia of Genes and Genomes (KEGG; Version 2021).

Pathway analysis of kinase activity in rd1 vs WT explants yielded as the major associated pathways Neurotrophin signaling pathway, proteoglycans in cancer, circadian entrainment, insulin resistance, HIF-1 signaling pathway, long-term potentiation, MAPK signaling (Fig. 4 a). After treatment of *rd1* explants with CN03, the high scoring pathways *i.e*. insulin resistance, mTOR, MAPK signaling, long-term potentiation, circadian entrainment, and HIF-signaling (Fig. 4 b) were the same as found in Fig 4a.

To identify the putative kinases involved in retinal degeneration based on rd1 model and PKG inhibitor CN03 treatment, we compared the kinases list derived from each comparison as represented in a Venn diagram (Fig. 4 c). Here, almost 77% of kinases with high activity in *rd1* as compared to WT were overlapping with the kinases with reduced activity in *rd1* retinal explants treated with CN03.

### PKG target validation for retinal localization

Next, we tested whether peptides that were differentially phosphorylated in the three retinal explant groups (*i.e*., WT, *rd1*, and *rd1*/CN03) were expressed in the retina. While some of the corresponding proteins had already previously been shown to be present in mouse retina (Table 2), we performed immunostaining on retinal tissue sections derived from P11 WT mice to assess the retinal expression and cellular localization for six proteins with high PKG preference (Table 2) and potentially connected to photoreceptor degeneration.

The analysis of the WT retinal sections immunostained for Cyclic AMP-responsive element-binding protein 1 (CREB1) showed a near ubiquitous distribution in all nuclear layers of the retina (Fig. 5). The potassium voltage-gated channel subfamily A, member 3 (K_v_1.3; KCNA3) was found to be localised in the ONL, possibly in photoreceptor axons, and in two discrete sublamina of the IPL (38), where the synapses between bipolar cell axons and ganglion cell dendrites reside. The potassium voltage-gated channel subfamily A, member 6 (Kv1.6; KCNA6) was expressed in the ganglion cell layer (GCL) and the nerve fibre layer (NFL). DNA topoisomerase 2-α (TOP2A) expression was restricted to the GCL. 6-phosphofructo-2-kinase/fructose-2,6-biphosphatase 3 (F263) – an enzyme involved in the control of glycolytic flux (39) appeared to be present in the OPL, likely in horizontal cells, as well as in the GCL. Finally, staining with the antibody directed against glutamate ionotropic receptor kainate 2 (GRIK2) confirmed its presence in the INL and GCL, in line with previous findings (31). Taken together, the immunostaining data confirm the retinal expression of several of the discovered PKG targets (24) (Fig. 5). The difference in phosphorylation of these targets between WT, *rd1* and *rd1*/CN03 retinas may suggest a potential role for these PKG substrates in the mechanism leading to photoreceptor cell death.

**Figure 5:**
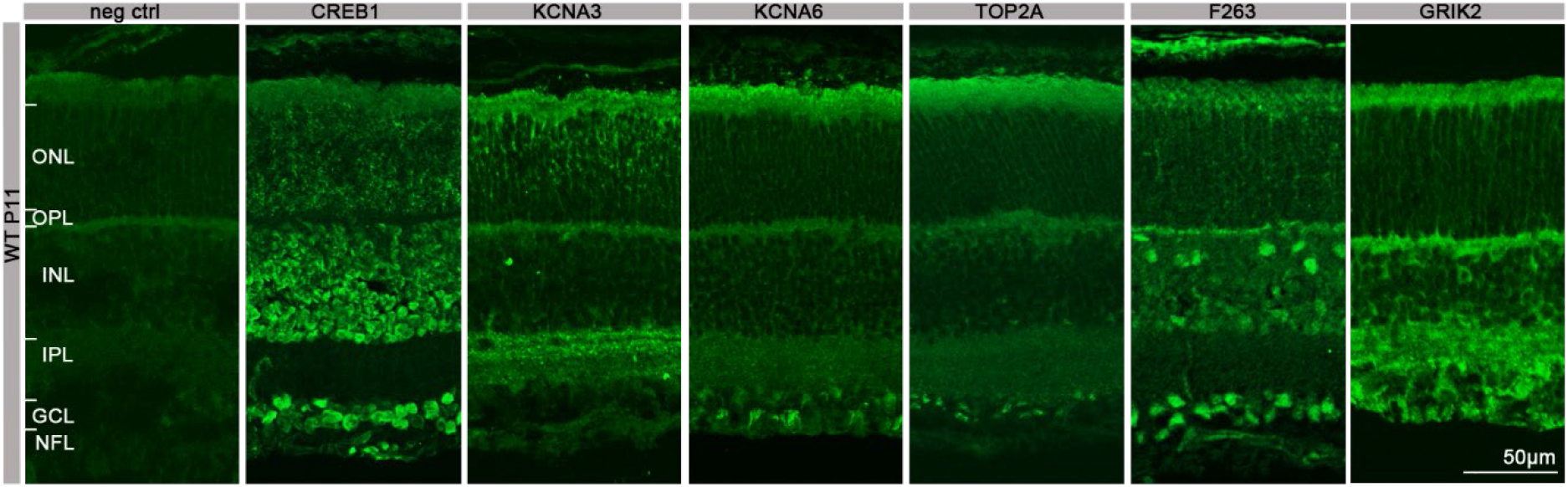
Expression and localisation of PKG target proteins in the retina. The panel shows retinal cross-sections derived from P11 WT mice and stained with secondary antibody for negative control (neg. ctrl.), anti-CREB1, anti-KCNA3, ant-KCNA6, anti-TOP2A, anti-F263, or anti-GRIK2. ONL=outer nuclear layer, OPL=outer plexiform layer, INL=inner nuclear layer, IPL=inner plexiform layer, GCL=ganglion cell layer, NFL=nerve fibre layer.

## Discussion

Excessive activity of PKG has been directly linked to photoreceptor cell death and its inhibition has been shown to provide photoreceptor protection in several *in vivo* IRD models (20,21). Yet, at present it is still unclear how PKG exerts its detrimental effects and what the protein targets are that, when phosphorylated by PKG, mediate photoreceptor cell death. Here, we combined PKG inhibitor treatment with multiplex peptide microarray technology and immunohistochemistry to identify novel PKG phosphorylation targets in the retina.

### PKG inhibition mediates photoreceptor neuroprotection

Over-activation of PKG has been connected to neuronal cell death in different experimental settings and conditions, including in human neuroblastoma derived cell cultures (40), peripheral nerve injury (41), and in photoreceptor degeneration (5,42). For photoreceptor degeneration an excessive accumulation of cGMP was already established in the 1970s and while it had already then become evident that high cGMP-levels were cytotoxic (43,44), it was unclear how cGMP would exert its negative effects. Research initially focussed on a proposed detrimental role of cyclic-nucleotide-gated (CNG) channels (45,46), however, in recent years it has become increasingly obvious that PKG-signalling plays a major role in photoreceptor degeneration (20,47). Our study, showing that the inhibitory cGMP analogue CN03 reduces photoreceptor cell death, is in line with earlier studies in which PKG inhibitors were found to preserve photoreceptor viability and functionality in a variety of models for inherited photoreceptor degeneration (20,21). Nevertheless, what protein targets of PKG exactly are responsible for its cell death promoting effects is unclear. Since PKG may phosphorylate hundreds of substrates it is important to identify those relevant for degeneration in a tissue and cell type specific context (48).

### PKG inhibition and peptide microarray technology for the prediction of PKG phosphorylation targets

To identify the biological pathways leading to photoreceptor cell death, the kinase activity of retinal samples was studied on the PamChip^®^ microarray platform, which allowed investigating the phosphorylation status of 142 peptides simultaneously. The *rd1* retinal explants showed an overall increased phosphorylation of peptides, indicating a higher kinase activity and the possible involvement of kinases from the AGC and CAMK families. Substrates to both isoforms of PKG; PKG1 and PKG2 showed higher activity in *rd1* retina as indicated by higher phosphorylation (Table 2). The specific inhibition of PKG activity with the inhibitor CN03 in organotypic retina explants allowed to profile kinase activity and response to PKG inhibition using PamChip^®^ technology. CN03 treatment showed a decrease in peptide phosphorylation with significantly reduced signal on seventeen peptides. Among the significantly regulated peptides, CREB1_126_38 has already been reported as PKG1 substrate (24). VASP, another well-known PKG substrate (21,24), was also affected, but ranked much lower on our substrate list (rank 22), suggesting that in a retinal context PKGs may prefer other substrates.

Recent studies identified additional cGMP-interacting proteins such as PKAs, PKCs, and CaMKs as upregulated in IRD retinas, which might be interesting in developing new therapeutic targets (49,50). These recent findings corresponded well with our study where we also identified these same kinases as activated in *rd1* diseased retinal explants and inhibited by the PKG inhibitor, CN03. The significant difference in phosphorylation between untreated *rd1* and CN03-treated *rd1* retinal explants indicates that PKG inhibition influenced phosphorylation of these peptides, whose corresponding proteins are therefore candidate PKG targets *in situ* and may play a role in the mechanism of photoreceptor cell death. We focused on those peptides for which PKG has a high preference, *i.e*., peptides with a high PKG score (Table 2), and confirmed their presence in the retina by either literature search or by additional immunofluorescence analysis.

### Novel targets for PKG in the neuroretina

One of the identified targets in our analysis was KCNA6, which, together with KCNA3 (rank 20 based on p value), belongs to the K_v_1 family of voltage-dependent potassium channels. The Kv1 family mainly consists of channels with a slow and delayed activation. Immunoreactivity studies revealed expression of both KCNA3 and KCNA6 in the mouse retina (29), which is in line with our immunofluorescence results. The Kv1 family is involved in the regulation of progression through cell cycle checkpoints by defining the membrane potential and ensuring the driving force for calcium and chloride entry. Surprisingly, these proliferation-related channels also appear to play a role in regulating cell death, making them interesting for our study (51). It has been shown that within a few minutes of apoptosis induction KCNA3 is inhibited by the cluster of differentiation 95 (CD95) *via* tyrosine phosphorylation (52). Other studies have shown the activation of KCNA3 at an early stage of apoptosis, as well as its contribution to a decrease in apoptotic volume, also known as ‘cell shrinkage’ in lymphocytes (53). In the retina, inhibition of KCNA1 and KCNA3 *in vivo* was protective for retina ganglion cells (RGCs) after optic nerve axotomy (54). The localization of KCNA6 and KCNA3 in the WT mouse retina and their different phosphorylation by PKG in *rd1* and CN03 treated *rd1* retinal explant cultures suggest a role in photoreceptor cell death. Past studies on the involvement of Kv1 channels in cell death, although conflicting, strengthen this idea (51–53). Establishing whether PKG-mediated phosphorylation of Kv1 channels involves activation or inhibition of these channels would help shed light on their contribution to photoreceptor death.

Among the various differentially phosphorylated substrates located in the retina of both *rd1* and WT, CREB1 is a known target of PKG (55,56). CREB1 is a transcription factor implicated in neuronal survival (57) and constitutively expressed in different types of human cancer (58). As shown by the *ex vivo* results, CREB1 is widely distributed in the mouse retina. Furthermore, through upstream kinase analysis we predicted the activation of several other kinases such as CaMK4, PKC theta, and PKA, known to target CREB (59–61). Thus, the abnormal activity of PKG as well as that of the other kinases can lead to increased CREB1 phosphorylation, in contrast to previous studies in which downregulation of CREB1 was associated with photoreceptor degeneration in both *rd1* and *rd10* mice (22,62). The increase in activity of kinases that phosphorylate CREB combined with an increase in CREB peptide phosphorylation in *rd1* may be due to the activation of parallel signals after the cellular insult: the induction of cell death and the activation of a CREB1-directed survival program (63).

Other PKG targets that we have identified and localized in the retina include vitronectin (VTNC) and F263. VTNC is a cell adhesion protein, upregulated in inflammation and traumatized tissues (64) and a major component of extracellular deposits specific to age-related macular degeneration (AMD) (65). In the same study, VTNC mRNA expression has been found distributed in the various layers of the human retina. Its high phosphorylation in *rd1* explants might be linked to increased inflammation, which is an early phenomenon observed for degenerative retinal disorders such as RP, AMD, and diabetic retinopathy (DR) (66). If so, VTNC would have the potential to be a predictive biomarker for certain retinal degenerative diseases.

6-phosphofructo-2-kinase/fructose-2,6-biphosphatase 3 or F263 is a pro-glycolytic enzyme. Its activation following excitotoxic stimulation has been associated with neuronal cell death (67). Furthermore, dysregulation of the HIF1-F263 pathway has been proposed as crucial for two key aspects of the pathogenesis of DR, namely angiogenesis and neurodegeneration (68). With *ex vivo* analysis we localized F263 in GCL and INL while it was not detected in ONL. The identification of F263 as a target of PKG and its high phosphorylation in *rd1* compared to WT suggests a change in retinal energy metabolism during degeneration and may also help to understand the mechanism of aerobic glycolysis in the retina, which remains relatively poorly understood to date (69).

### Metabolic pathways in the retina regulated by PKG

The pathway analysis showed insulin resistance and mTOR as the top pathways inhibited by CN03. mTOR is a serine/threonine kinase which regulates protein synthesis, cellular metabolism and autophagy (70). The mTOR pathway regulates neurogenesis in the eye and is crucial to normal development of retina and the optic nerve (71). Activated mTOR signaling has been shown to be involved in retinal neurodegenerative diseases such as DR and AMD (72,73). Treatment with the mTOR inhibitor rapamycin, improved mitochondrial dysfunction and provided neuroprotection to 661W cells, a cellular model that share certain features with photoreceptors (74). Similarly, inhibition of the mTOR/PARP-1 axis leads to photoreceptor protection against light-induced photoreceptor cell death (70). However the targeting of this pathway axis in IRD as treatment strategy is still contentious as there are also studies that show mTOR stimulation delays cone cell death in IRD (75,76).

Among the peptides whose phosphorylation was high in untreated *rd1* retinal explants and significantly decreased in CN03-treated *rd1* retinal explants was F263, a key regulator of glycolysis (77). Photoreceptors metabolize glucose through aerobic glycolysis (the ‘Warburg effect’) to satisfy their energy demand for the maintenance of the dark current, as well as the recycling of visual pigments and the renewal of photoreceptor OS (78,79). As a consequence of the high energy demand, the retina has an elevated oxygen consumption, probably the highest in the body, and is therefore particularly prone to oxidative stress and reactive oxygen species (ROS)-induced mitochondrial damage (80). The high phosphorylation of F263 found in the *rd1* retina may be linked to an abnormal increase in metabolic activity in the retina resulting in mitochondrial damage. Considering that mitochondrial dysfunction has been associated with RP (80), the identification of F263 as a substrate of PKG and its possible role in the degeneration process could help to clarify the metabolic state of the diseased retina.

## Conclusion

Using multiplex peptide kinase array technology, we provide further evidence for the likely involvement of PKG in degenerating *rd1* photoreceptors *in vitro* and confirmed the already known neuroprotective effects of the PKG inhibitor CN03 (21). Furthermore, we identified several novel downstream PKG targets that might play a role in cGMP/PKG-mediated photoreceptor degeneration. This will form the basis for future studies which may further elucidate the role of PKG target phosphorylation as well as the beneficial effects of CN03 and could, furthermore, determine their relevance as novel diagnostic and therapeutic biomarkers for retinal degenerative diseases. For example, the use of phospho-specific antibodies may allow to confirm a reduction of target phosphorylation and, in *in situ* studies, allow to localize in which cellular compartment it occurred. Finally, our results further connect several metabolic pathways with retinal degeneration, including insulin, mTOR, and Hif1 signaling, which in future studies may help to understand the complex mechanisms behind photoreceptor cell death.

## Materials and Methods

### Animals

C3H Pde6b^*rd1*/*rd1*^ (*rd1*) and congenic C3H wild-type (WT) mice were housed under standard light conditions, had free access to food and water, and were used irrespective of gender. All procedures were performed in accordance with the law on animal protection issued by the German Federal Government (Tierschutzgesetz) and approved by the institutional animal welfare office of the University of Tübingen.

#### cGMP analogues synthesis

Synthesis of the cyclic nucleotide analogue CN03 was performed by Biolog Life Science Institute GmbH & Co. KG according to previously described methods (21) (https://patentscope.wipo.int/search/en/detail.jsf?docId=WO2018010965)

### Organotypic retinal explant cultures

Preparation of organotypic retinal cultures derived from *rd1* (n = 10) and C3H (n= 5) animals was performed as described previously (26,81). Animals were sacrificed at postnatal day (P)5. Eyes were rapidly enucleated and incubated in R16 serum-free, antibiotic-free culture medium (07491252A; Gibco) with 0.12% proteinase K (21935025; ICN Biomedicals Inc.) for 15 min at 37°C. Subsequently, eyes were incubated in 20% foetal bovine serum (FCS) (F7524; Sigma) in order to block proteinase K activity. This step was followed by rinsing in R16 medium. Under a laminar-flow hood and sterile conditions, the anterior segment, lens, vitreous, sclera, and choroids were removed from the eyes. The retina with the RPE still attached was cut in four points resembling a four-leaf clover and transferred to a culture membrane insert (3412; Corning Life Sciences) in a six-well culture plates with completed R16 medium with supplements (26). The retinal explants were incubated at 37°C in a humidified 5% CO2 incubator and left undisturbed for 48h. At P7 and P9 medium was changed every second day *i.e*. with replacement of the full volume of the complete R16 medium, 1mL per dish, with fresh medium. In this context, retinal explants were either treated with CN03 at 50 μM (dissolved in water) or kept as untreated control. For retinal explant lysis, culturing was stopped at P11, retinal explants were snap frozen in liquid nitrogen and stored at −80 °C. For retinal explants cross-sectioning preparation, culturing was stopped at P11 by 45 min fixation in 4% paraformaldehyde (PFA), cryoprotected with graded sucrose solutions containing 10, 20, and 30% sucrose, embedded in optimal cutting temperature compound (Tissue-Tek) and then cut into 12μm sections.

### TUNEL assay

Representative results showing the protective effects of CN03 on photoreceptor cell death at a concentration of 50 μM, were obtained using terminal deoxynucleotidyl transferase dUTP nick end labelling (TUNEL) assay (82) (based on in Situ Cell Death Detection Kit, 11684795910, red fluorescence; Sigma-Aldrich) on sections derived from *rd1* and C3H retinal explant cultures. DAPI (Vectashield Antifade Mounting Medium with DAPI; Vector Laboratories) was used as blue fluorescence nuclear counterstain. Images were captured using 7 Z-stacks with maximum intensity projection (MIP) on a Zeiss Axio Imager Z1 ApoTome Microscope MRm digital camera (Zeiss, Oberkochen, Germany) with a 20x APOCHROMAT objective. The excitation (*λ_Exc_*) / emission (*λ_Em_*.) characteristics of the filter sets used for the fluorophores were as follows (in nm): DAPI (*λ_Exc_*. = 369 nm, *λ_Em_*. = 465 nm) and TMR red (*λ_Exc_* = 562 nm, *λ_Em_* = 640 *nm*). Adobe Photoshop (CS5Adobe Systems Incorporated, San Jose, CA) was used for image processing.

### Materials for retina explant lysis

Mammalian protein extraction reagent (M-PER™), Halt™ protease and phosphatase inhibitor cocktails and the Coomassie Plus (Bradford Assay) kit were purchased from Thermo Fischer Scientific.

### Retinal Explant Lysis

The retinal explant samples were lysed with lysis buffer (MPER with 1:100 phosphatase inhibitor cocktail and protease inhibitor cocktail reagents) for 30 mins on ice. The lysate was centrifuged at 16 000 x *g* for 15 min at 4 °C. The supernatant was immediately aliquoted, flash frozen, and stored at −80 °C. The protein content of the lysate was measured using the Bradford Protein Assay (83).

### Kinase activity measurements

The kinase activity for the retina lysates was determined on STK PamChip^®^ with four arrays, each comprising of 142 peptides derived from the human phosphoproteome, according to the instructions of the manufacturer (PamGene International B.V., ‘s-hertogenbosch, North Brabant, The Netherlands). The peptide names consist of the protein they are derived from and the first and last amino acid positions in that protein. The phosphorylated Serine/Threonine amino acid residues are detected by a primary antibody mix, which is then confirmed by addition of FITC-conjugated secondary antibody (84). The assay mix consisted of protein kinase buffer (PamGene International BV. ‘s-Hertogenbosch, North Brabant, The Netherlands), 0.01% BSA, STK primary antibody mix, ATP (400 μM) and retina tissue lysate (0.25 μg/array).

### Instrumentation for kinase activity measurements

All the experiments were performed on PamStation12^®^ where up to 12 assays can be performed simultaneously (PamGene International B.V., ‘s-Hertogenbosch, North Brabant, The Netherlands). To prevent unspecific antibody binding, the PamChips^®^ were first blocked with 2% BSA, by pumping it up and down 30 times through the arrays. The chips were then washed three times with Protein Kinase Buffer and assay mix was applied. The assay mix was pumped up and down through the arrays for 60 mins. Afterwards, the arrays were washed and FITC labelled secondary antibody mix was applied on the arrays. The images of the arrays were recorded at multiple exposure times (85).

### Data analysis

The signal intensity of each peptide spot on the array for each time point was quantified by BioNavigator^®^ software version 6.3.67.0 (PamGene International B.V., ‘s-Hertogenbosch, North Brabant, The Netherlands). For each spot, the signal intensity at the different exposure times was combined to a single value by exposure time scaling (85). The resulting values were log2 transformed and the overall differences in STK profile between *rd1* NT *vs*. WT or *rd1* CN03 *vs*. *rd1* NT were visualized as Heatmaps and Violin Plots which were generated in R software ((R version 4.0.2, 2020 The R Foundation for Statistical Computing). For heatmaps, hierarchal clustering of peptides was performed using the average-linkage method and clustering was shown as dendrograms on the y-axis of heatmaps. Unpaired *t*-tests were used to determine significant differences (*p* < 0.05) in phosphorylation intensity between the groups. Results were represented as Volcano Plots (GraphPad Prism version 9.2.0).

### Upstream kinase analysis

Information on kinases that could be responsible for peptide phosphorylation differences between the two sample groups was obtained through the STK Upstream Kinase Analysis tool of BioNavigator^®^ (85). This software integrates known interactions between kinases and the phosphorylation sites as provided in databases such as HPRD, PhosphoELM, PhosphositePLUS, Reactome, UNIPROT and predicts the differentially active kinases between the groups. The results of this analysis are described by two parameters: The Kinase Statistics, indicating the size and direction of the change for each kinase and the Kinase Score, ranking the kinases by the likelihood of this kinase being involved.

The Kinase Statistic depicts the overall change of the peptide set that represents a kinase. For instance, a larger positive value indicates a larger kinase activity in either *rd1* explants in comparison to WT or CN03 treated explants in comparison to untreated explants. The Kinase Score is the result of two permutation analyses. The Kinase Score is calculated by addition of the Significance Score and the Specificity Score. The Significance Score indicates the significance of the change represented by the Kinase Statistic between two groups (using 500 permutations across sample labels). The Specificity Score indicates the specificity of the Kinase Statistic with respect to the number of peptides used for predicting the corresponding kinase (using 500 permutations across target peptides). The kinases are ranked by the Kinase Score. The highest ranking predicted kinases from the significant STK peptide sets are represented on a phylogenetic tree of the human protein kinase family generated in Coral, a web-based application http://phanstiel-lab.med.unc.edu/CORAL/) (86).

### Pathway analysis

Pathway analysis was performed in Enrichr (https://maayanlab.cloud/Enrichr/), which is a comprehensive resource for curated gene sets and a search engine that accumulates biological knowledge for further biological discoveries (36,37). We used differentially phosphorylated peptides and predicted upstream kinases as input list for Erichr analysis. Visualization was performed in GraphPad Prism (version 9) using known matrices from the analysis.

### Histology

For retinal cross-section preparation, the eyes were marked nasally and cornea, iris, lens, and vitreous were carefully removed. The remaining eyecups were fixed in 4% PFA for 2 h at room temperature. Incubation with graded sucrose solutions was performed for cryoprotection. Eyes were embedded in Tissue- Tek and cut into 14 μm sections. Immunostaining was performed by incubating with primary antibody against rabbit CREB1 (1:200; Proteintech), KCNA3 (1:200; Alomone labs), KCNA6 (1:300; Alomone labs), F263 (1:100; Abcam), TOP2A (1:500; Proteintech), GRIK2 (1:100; Invitrogen), at 4 °C overnight. Alexa Fluor 488 antibody was used as secondary antibody. Sections were mounted with DAPI. Images were captured using 9 Z-stacks with maximum intensity projection (MIP) on a Zeiss Axio Imager Z1 ApoTome Microscope MRm digital camera (Zeiss, Oberkochen, Germany) with a 20x APOCHROMAT objective. The excitation (*λ_Exc_*) / emission (*λ_Em_*.) characteristics of the filter sets used for the fluorophores were as follows (in nm): DAPI (*λ_Exc_*. = 369 nm, *λ_Em_*. = 465 nm) and AF488 (*λ_Exc_* = 490 nm, *λ_Em_* = 525 nm). Adobe Photoshop (CS5Adobe Systems Incorporated, San Jose, CA) was used for image processing.

## Acknowledgements

We thank Norman Rieger for excellent technical assistance and Rik de Wijn, Savithri Rangarajan and Faris Naji for helpful discussion on data analysis. This research was funded by the European Union Horizon 2020 Research and Innovation Programme- *trans*Med under the Marie Curie grant agreement No. 765441 [(*trans*Med; H2020-MSCA-765441)].

## Supplementary Figures

**Figure S1.**
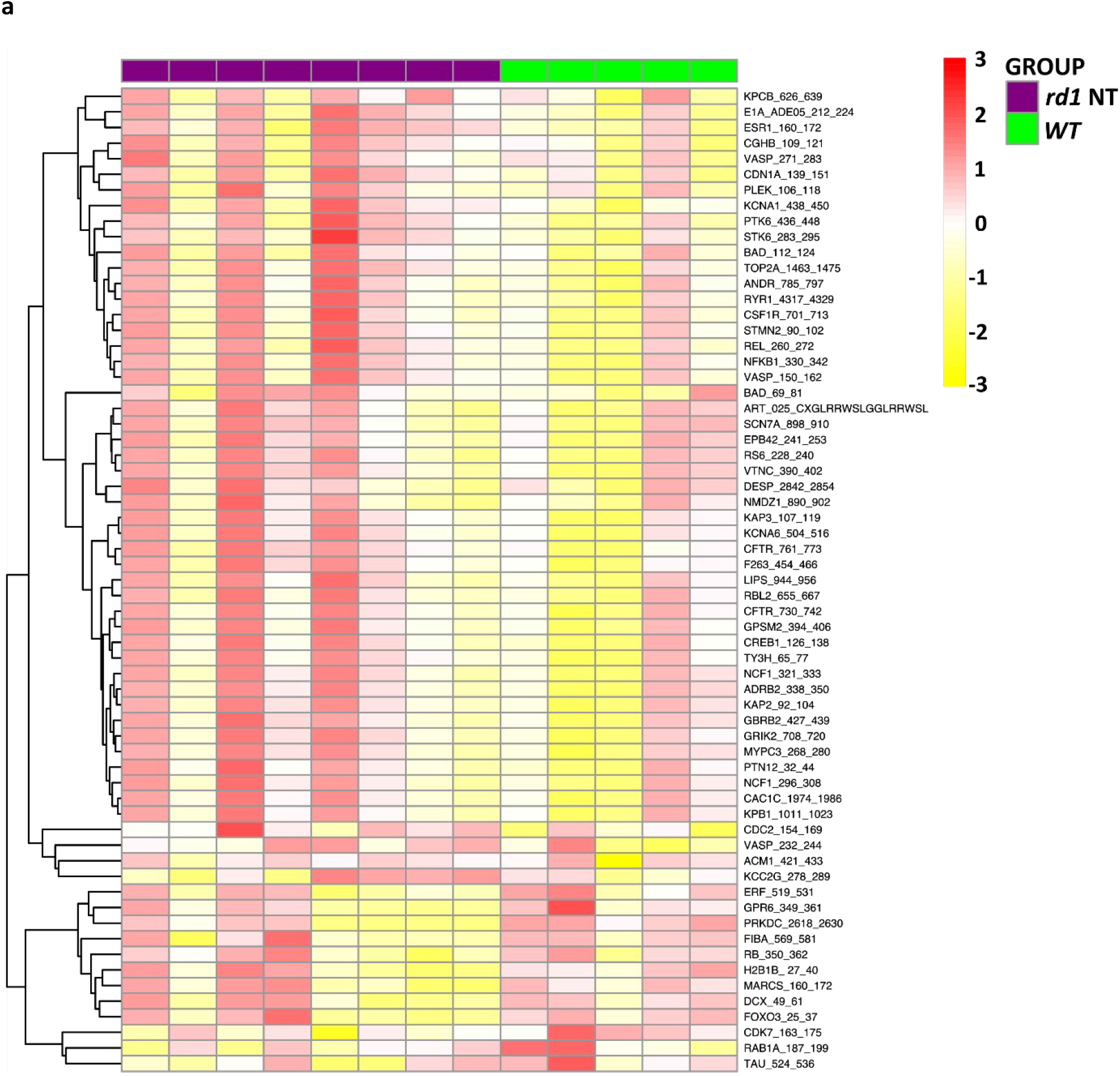
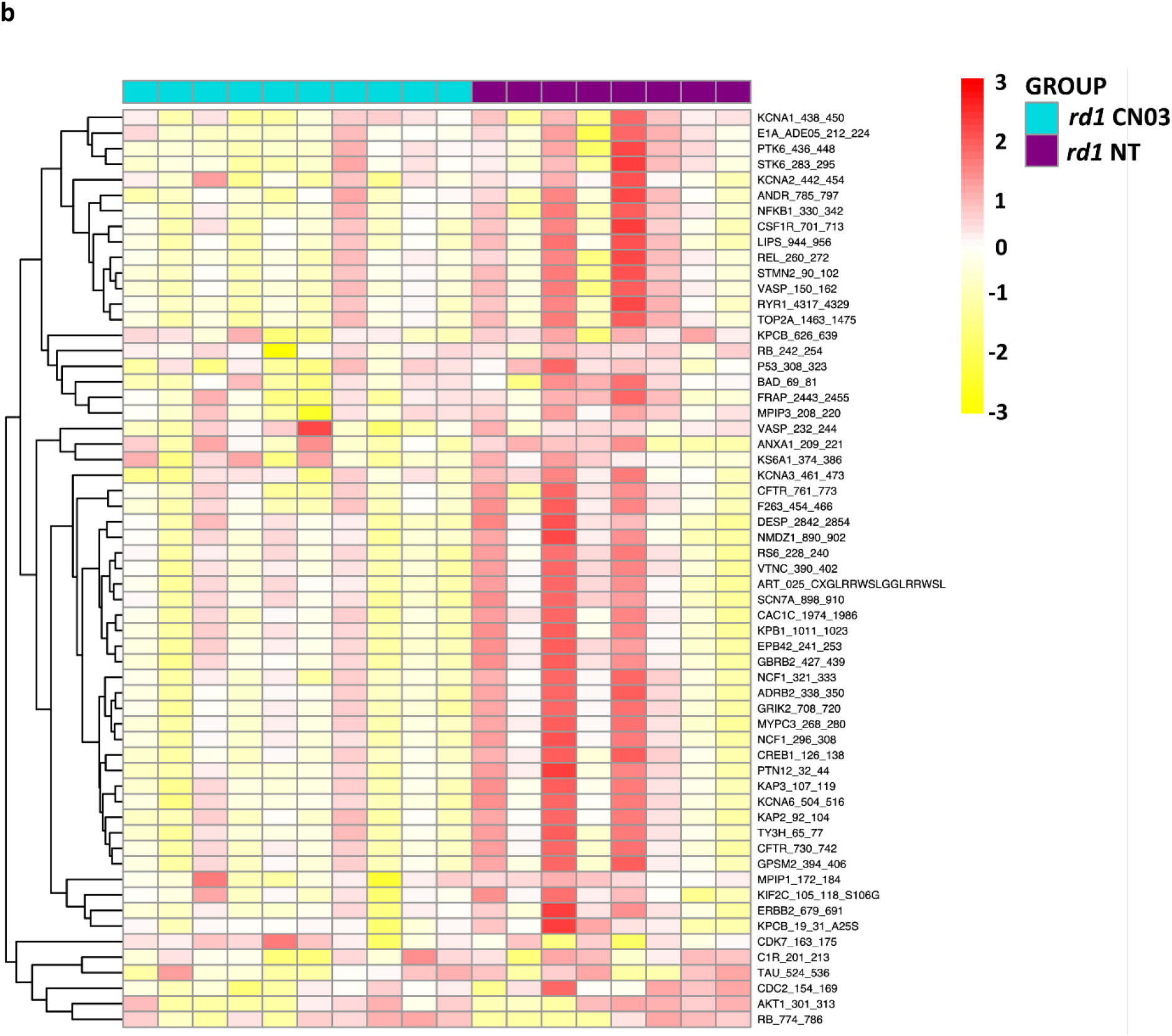
Heatmap representing the overall serine/threonine kinase activity in retinal explants. **a**) Comparison of protein phosphorylation between *rd1* vs. WT retinal explant cultures. **b**) comparison of CN03 treated *vs*. non-treated (NT) *rd1* retinal explant cultures. The phosphorylated peptides are clustered hierarchically as explained in ‘Data Analysis’ section (red=high phosphorylation; yellow=low phosphorylation).

